# Dsg2 truncation causes a lethal barrier breakdown in mice

**DOI:** 10.1101/2024.07.01.601522

**Authors:** Daniela Kugelmann, Elisabeth S. Butz, Alexander Garcia Ponce, Mariya Y. Radeva, Jessica Neubauer, Matthias Hiermaier, Christian Siadjeu, Natalie Burkard, Desalegn Egu, Anja K. E. Horn, Christoph Schmitz, Katri S. Vuopala, Outi Kuismin, Nicolas Schlegel, Jens Waschke

**Affiliations:** Chair of Vegetative Anatomy, Institute of Anatomy, Faculty of Medicine, LMU Munich, Munich, Germany; Systematics, Biodiversity and Evolution of Plants, Faculty of Biology, LMU Munich, Munich, Germany; Department of General, Visceral, Transplant, Vascular and Pediatric Surgery University Hospital Wuerzburg, Wuerzburg, Germany; Chair of Neuroanatomy, Institute of Anatomy, Faculty of Medicine, LMU Munich, Munich, Germany; Department of Pathology, Lapland Central Hospital, Rovaniemi, Finland; Department of Clinical Genetics, Oulu University Hospital, OULU, Finland

## Abstract

Inflammatory bowel diseases (IBD) such as Crohn’s disease (CD) have a complex aetiology with alterations of both the intestinal epithelial barrier and the IL23/IL17 immune response. Here, we investigated the role of a novel mutation in the desmosomal cadherin desmoglein 2 gene (*DSG2*) in the pathogenesis of IBD. DSG2 is known to regulate intestinal epithelial barrier integrity. Genetic analysis of a CD patient revealed a novel likely pathogenic *DSG2* mutation leading to a truncated protein lacking part of the intracellular domain. We generated an enterocyte-specific mouse model, recapitulating the human mutation to study how the cytoplasmic truncation of *Dsg2* affects intestinal barrier properties systemically. Moreover, we analysed the intestinal genetic profile in these mice and compared it to IBD patients. We describe a first CD patient with a rare mutation in the *DSG2* gene causing cytoplasmic truncation with affects Dsg2 mobility. Mice with enterocyte-specific Dsg2 truncation suffered from a lethal intestinal barrier defect and presented a skewed IL17 response similar to CD patients. We identified the desmosomal cadherin Dsg2 as a regulator of the skewed IL17 response. These data indicate that desmosomes regulate inflammation similar to psoriasis which explains why the same novel immune therapies are effective for both diseases.

## Introduction

Inflammatory bowel diseases (IBD) such as Crohn’s disease (CD) and ulcerative colitis (UC) are characterized by a complex pathogenesis which is not completely understood. CD can affect all parts of the gastrointestinal tract whereas UC mainly affects the colon^1^. The multifactorial pathogenesis involves genetic predisposition, composition of gut microbiota, immune system dysfunction and environmental factors as triggers for IBD^2 3^. Genome-wide association studies identified more than 200 risk gene loci^4^. It is well established that a skewed IL23/IL17 immune response significantly contributes to IBD pathogenesis which is targeted by novel immune therapies^5–8^. Another hallmark of IBD is intestinal epithelial barrier dysfunction^3^. The intestinal barrier consists of the single-layered intestinal epithelium which under normal conditions is sealed by the apical junctional complex consisting of tight junctions (TJ), adherens junctions (AJ) and desmosomes^9^. Based on the observation that the desmosomal adhesion molecule desmoglein (Dsg) 2 is required for intestinal epithelial barrier function under basal conditions, the role of desmosomes in the pathogenesis of IBD was investigated^3^. Indeed, Dsg2 and also desmocollin (Dsc) 2 were consistently shown to be reduced in biopsies taken from patients with CD and UC^10–12^ where the ultrastructure of desmosomes but not AJ was found to be altered^13^. Mouse models revealed that loss of Dsg2 or Dsc2 causes intestinal barrier disruption and compromises intestinal mucosal repair^12–14^. All these data demonstrate that desmosomal adhesion is disturbed in IBD and may constitute a gain-of-function toxicity. However, the functional interplay between altered intestinal epithelial barrier properties and dysregulation of the immune system is unclear at present. Here, we report a first IBD patient with a mutation in the *DSG2* gene leading to a truncation of the protein within the cytoplasmic domain. We demonstrate that the mutation severely affects desmosome turnover and causes a lethal intestinal epithelial barrier loss in neonatal mice paralleled by a skewed IL17 response comparable to CD patients.

## Methods

### Consent and study information

Written informed consent was obtained from the patient and the study project on Rare Genetic Diseases in Northern Finland was approved by the Ethics Committee of North Ostrobothnia’s Hospital District. During colonoscopy biopsy samples were taken from the colon and used for morphological analysis. Biopsies from unaffected intestinal regions from the patient served as control.

### Genetic analysis and Clinical Information

Whole exome sequencing of the patient was ordered from the clinical diagnostic laboratory Centrogene Rostock, Germany.

### Cell culture

DLD1 cells defective for Dsg2 and Dsc2 (DLD-1 ΔDsc2 ΔDsg2) were cultured in Dulbecco’s modified Eagle medium (Life Technologies, Carlsbad, CA, USA) supplemented by 10 % fetal bovine serum (Biochrom, Berlin, Germany), 50 U/mL penicillin, and 50 U/mL streptomycin (both AppliChem, Darmstadt, Germany), and cultivated in a humidified atmosphere containing 5 % CO_2_ at 37 °C. DLD1 knockout cells were generated in the lab of Suzuki (Kwansei Gakuin University, Japan)^15^ using the CRISPR/Cas9 system with the pX330 vector (Addgene, Cambridge, MA, USA) according to the method published by Cong et al.^16^ as described previously^17^.

### Plasmids and transfection

DLD-1 ΔDsc2 ΔDsg2 enterocytes were transfected with either DSG2-EGFP, DSG2Leu772*-EGFP or the control vector pEGFP-EV using 4D-Nucleofector (Lonza, Basel, Switzerland) according to the manufactureŕs instructions. For the electroporation 80 to 95 % confluent DLD-1-cells were trypsinized and counted. An amount of 10^6^ cells were resuspended in 100 µl of SE Cell Line Solution (Cat. No. V4XC-2024, Lonza, Basel, Switzerland) containing 3.5 µg of the respective plasmid and electroprated with pulse DG113. Experiments were performed 24 hours after electroporation.

The DSG2-EGFP plasmid encoding for C-terminally eGFP-tagged human wild type Dsg2^18^ was kindly provided by Katja Gehmlich (BHF Centre of Research Excellence Oxford, University of Oxford, Oxford, UK). DSG2Leu772*-EGFP encodes for the latter DSG2 bearing a stop codon after amino acid leucine at position 772.

### Fluorescence Recovery After Photobleaching (FRAP)

FRAP studies were performed as described previously^19 20^. For the measurement cells were transfected with either DSG2-EGFP or DSG2Leu772*-EGFP) and thereafter seeded in 8-well imaging chambers (Ibidi, Martinsried, Germany). The measurements were performed on a SP5 inverted confocal microscope with a 63x HC PL APO NA=1.2 objective (Leica, Wetzlar, Germany) at 37 °C with 5 % CO_2_ and constant humidity using a cage incubator (OKOLAB, Burlingame, CA, USA) and analyzed with the FRAP wizard software (Leica, Wetzlar, Germany). The regions of interest were defined along the cell membranes of two neighboring transfected cells. After five frames of recording the pre-bleach intensity, the EGFP signal was bleached using the 488-nm laser line at 100 % transmission for 10 frames and recovery of the fluorescence was recorded for an appropriate amount of time with varying duration of frames. The fraction of mobile molecules was determined by:

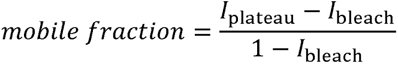

Here, *I*_plateau_ is the plateau intensity which is reached after sufficient recovery time and *I*_bleach_ is the minimal intensity that was achieved right after bleaching. Both intensities were normalized to the pre-bleach intensity.

### Western blot analysis

For lysates of intestinal segments, the entire intestine was flushed with PBS. Terminal ileum and colon were extracted and used for lysates. Tissue was collected in sodium dodecyl sulfate (SDS)-containing lysis buffer supplemented with protease inhibitors (cOmplete, Cat. No.: 11697498001, Roche, Basel, Switzerland) and homogenized using gentleMACS^TM^ Octo dissociator (Miltenyi, Bergisch Gladbach, Germany) according to the manufacturer’s recommendation. Lysates were sonicated, protein concentration was determined using Pierce^TM^ BCA Protein Assay Kit (Cat. No. 23225, Thermo Scientific, Waltham, MA, USA) and equal protein amounts were mixed with SDS-containing Laemmli buffer and loaded onto SDS-polyacrylamide gels. After electrophoresis, proteins were transferred to nitrocellulose membranes, blocked in 5 % low-fat milk in Tris-buffer saline (TBS) with 0.0475 % Tween-20 (TBS-T) and subsequently incubated with specific primary antibodies diluted in 5 % milk powder or 5 % bovine serum albumin (BSA) in TBS-T at 4 °C overnight. Primary antibodies were used as follows: anti-Dsg2 (abx232544, 1:1000, Abbexa, Cambridge, UK), anti-claudin-2 (21H12) (Cat. No. 32-5600, 1:1000, Thermo Scientific, Waltham, MA, USA), anti-E-Cadherin (Cat. No. 610181, 1:1000, BD Biosciences, Heidelberg, Gemany), anti-Gapdh (OAF04737, 1:5000, Aviva Systems Biology, San Diego, CA, USA), anti-alpha-Tubulin (ab7291, 1:10^4^, Abcam, Cambridge, UK). The next day, membranes were incubated with horseradish-peroxidase-coupled secondary antibodies and protein bands were detected with enhanced chemiluminescence solution. Protein levels in the Western blots were quantified by densitometry using ImageJ software (National Institutes of Health (NIH), Bethseda, MA, USA).

### Immunohistochemistry- and Immunfluorescence analyis

For histologic analysis, the intestine was removed from the animal, washed in PBS and immediately fixed in 2 % paraformaldehyde (PFA) in PBS for 2 hours. Defined intestinal segments from jejunum, proximal and terminal ileum, ascending colon and rectum were then embedded in paraffin cut into 7µm thick sections. For hematoxylin and eosin stainings, sections were deparaffinized, washed with water and incubated with hematoxylin followed by eosin incubation. Sections were subsequently embedded in Xylol and DPX. Brightfiled images of the specimen were captured using a slide scanner (Mirax MIDI, Zeiss, Jena, Germany), equipped with a Stringray camera with a Plan-Apochromat objective (20x, Zeiss, Jena, Germany) and evaluated with the free software Case Viewer (3DHistech, Budapest, Hungary). For immunohistochemistry, paraffin-embedded sections from human biopsy speicmen and murine intestine were deparaffinized and boiled in EDTA-TRIS buffer for antigen retrieval. Sections were then permeabilized in 0.3 % Triton/PBS for 30 min, blocked in 5 % BSA/ NGS/TBS for 60 min, and incubated in primary antibody diluted in BSA/NGS in TBS under humid conditions at 4 °C overnight. The study was performed using antibodies against Dsg2 (Cat. No. 610121, 1:100, Progen, Heidelberg, Germany, or Cat. No. BM5016, 1:20, Origene, Rockville, MD, USA), E-Cadherin (Cat. No. 610181, 1:100, BD Biosciences, Heidelberg, Gemany), Occludin (Cat. No. 40-4700, 1:100, Thermo Scientific, Waltham, MA, USA), Claudin 2 (Cat. No. 51-6100, 1:100, Thermo Scientific, Waltham, MA, USA). Next day, sections were washed and further incubated with fluorphore-tagged secondary antibodies (Dianova, Hamburg, Germany) for 1 h, stained with DAPI for 15 min, washed rigorously and mounted with n-propyl gallate (NPG)-containing mounting medium. Images were acquired with Leica confocal (SP5, Leica, wetzlar, Germany). Images were processed using Adobe Photoshop (Adobe Inc, San Jose, CA, USA).

### Intestinal barrier permeability ex vivo assay

The permeability of intestinal epithelial barrier of Dsg2_MUT_ and wildtype mice was assessed by an assay adopted from Mateer et al.^21^ (2016). Specifically, neonatal mice at postnatal day 7 were decapitated and the peritoneum was opened. The gastrointestinal tract was removed immediately, transferred in a PBS containing petri dish and cleaned from mesentery. A fragment of the distal small intestine was cut extracted revealing a free segment with two open ends. The tip of a feeding tube adjusted to a PBS-containing 1 mL syringe was plugged into the proximal end and closely tightened with a suture loop. The intestinal segment was flushed with 1 ml PBS, before the syringe was exchanged by a syringe filled with Fluoresceinisothiocyanat (FITC)-dextran MW 4,000 Da (Cat. No. 46944, Sigma Aldrich, Munich, Germany) dye, which was diluted in PBS to a final concentration of 2 mg/mL. Intestine was gently flushed with dye and the distal opening was closed with a suture-loop to obtain an approximately 3 cm long segment of gut (“intestinal sac”). The tissue was immediately transferred to a 25 ml baker containing 7 mL of pre-warmed, oxygenized phenol-red free DMEM medium (Cat. No. 21063029, Thermo Fisher Scientific, Waltham, MA, USA) and transferred into a water bath heated to 37 °C for the course of the experiment. 200 µl samples of medium were taken immediately after the specimen was put into the baker and 200 µl samples were consecutively collected every 5 minutes for a time period of 20 minutes. Oxygenized, heated pure medium was used as reference. To assess barrier permeability in heterozygous mice, 14 days old mice were investigated. Intestinal segments were prepared as described before, except, that intestinal sacs were filled with 4 kDa FITC solution alone or 4 kDa FITC solution supplemented with tumor necrosis factor α (recombinant murine TNFα, Peprotech, #315-01A, final concentration: 100 ng/mL), interleukin 1b (recombinant murine Il-1b, Sigma-Aldrich, #I5271, 10 ng/mL) and interferone γ (recombinant mouse INFγ, Cell Signaling, #39127 10 ng/mL) (cytomix challenge) and 200 µl samples were taken every 15 min for 1 hour, followed by every 30 min for a total of 4 hours. Samples were taken in duplicates and collected in a 96-well plate for detection. Fluorescent intensity of the medium as a measurement of the flux of the dye across the intestinal epithelial barrier was determined using a microplate reader (Infinite 200 Pro, Tecan, Männedorf, Switzerland) and appropriate software (i-contol^TM^, Tecan, Männedorf, Switzerland). Leakiness of the gut tissue over time and apparent permeability was subsequently analyzed using Microsoft Excel (Microsoft Office, Redmond, WA, USA).

### Mouse housing and collection

All animals were kept under conditions according to regulations and permission of the Government of Upper Bavaria, (Az. 55.2-2532.Vet 02-19-172), Germany. All experiments were performed in accordance with the relevant guidelines and regulations. Neonatal mice were collected between postnatal day 6 and 13 blinded to genotype. Afterwards, genotyping was performed on tail biopsies from mouse carcasses. On the day of mouse collection, size and total body weight of neonatal pups were determined, animals were terminated by decapitation and tissue from gastrointestinal tract was collected as follows: For protein lysates, intestine was removed, washed in PBS and up to 100 mg of tissue was collected in SDS-containing lysis buffer. To remove faeces and mucus the intestine was cut longitudinally and flushed with PBS. For immunohistochemistry, mouse tissue was washed in PBS and transferred to 2 % PFA. After 2 hours incubation at 4 °C, PFA was exchanged by PBS and the tissue was processed for paraffin sections.

### Generation of mice

An enterocyte-specific knock-in mouse model was established causing cytoplasmic truncation of Dsg2 in exon 14 (Dsg2_MUT_), mimicking the identified human mutation.

### Generation of conditional transgenic mouse for Dsg2 gene carrying nonsense mutation at the 3’ of exon 14 (Ex14)

The transition gene (g.) 2315T*>*G located into the terminal part of exon 14 (Ex 14) from human DSG2 gene was introduced, leading to substitution of protein (p.) Leucine (Leu) 772*>*premature stop codon (PSC) (i.e. DSG2Leu772/X). This location is specific, because an earlier stop (further up in Ex 14, or in one of the earlier exons) would indeed cause a termination of the message, but also impair the stability of the mRNA by Nonsense-Mediated Decay (NMD). According to the currently proposed mechanism for NMD, a stop codon localized 50-55 nucleotides upstream of the exon-exon junction is recognized as a PSC and once the NMD is stimulated, the targeted/irregular transcript is degraded by 5’ and 3’ exonucleases. As a result, the protein would not only be truncated, but the truncated protein will be also very poorly expressed. A notable exception is a stop that is closer than 50 nucleotides from the splice donor signal, as it is the particular case here. In the case of DSG2Leu772/X, a truncated peptide is expressed at physiological levels and this model must replicate this feature.

Similarly, to the reported in human mutation, where Leu at position 772 was exchanged into a PSC, a mouse strain was established carrying a PSC closed to the 3’ end of Ex 14 of mouse DSG2 gene. The latter, in analogy to the human gene, consists of 15 exons in total. In addition, mouse Dsg2 protein shares 75 % amino acid identity with the human Dsg2 as the protein sequence is convincingly homologous in the region where the mutation of interest is detected. Thus, a model expressing a truncated form of Dsg2 protein was developed, where due to the newly introduced PSC, the 3’ end of Ex 14 and the complete Ex 15 were not further translated into a native protein.

In brief, to generate the mutant in a conditional manner, an elegant targeting strategy was developed (figure 3A). Here, Ex 14 was subdivided into two partial exons (Ex 14A and Ex 14B) by inserting an artificial intron (intron 4 of mouse Dsg2, 132 bp plus a loxP site) right after the mutated site. Thus, the major part of Ex 14, i.e. Ex 14A (316 bp) was followed by an artificial intron and the first LoxP site used for excision by tissue specific Cre recombinase. The modified DNA fragment continue with a sequence representing the rest of the Ex 14 (mini Ex 14B; 14 bp in size) and the second LoxP site, which was incorporated into the intron originally positioned between Ex 14 and Ex 15. As a result of proper splicing, a native continual, full wild type (WT) mRNA was generated, which in turn was translated into the original protein. However, after excising Ex 14B by using tissue specific Cre recombinase, Ex 14A and Ex 15 fused and a stop site was activated resulting in production of a truncated form of the Dsg2 protein missing the amino acid sequence corresponding to the terminal end of Ex 14 and the complete Ex 15.

A standard approach based on a homologous recombination in embryonic stem (ES) cells was used for generation of Dsg2 conditional knock-in with PSC introduced at the terminal 3’ part of Ex 14 (supplemental figure 1). The method is based on a targeting vector with pBluescript KS-backbone, which carries two homologous regions enhancing successful recombination (supplemental figure 2). The subcloned 5’ homologous short arm (SA) contains 2,72 kb of randomly selected piece of genomic DNA (gDNA) surrounding the region of interest, while a fragment of 4,87 kb gDNA serves as a the 3’ recombination long arm (LA). The homology arms were generated by PCR using C57Bl/6N BAC DNA (clone RP23-93P4) as template. SA and LA are flanking a Neo cassette, inserted in between FRT sites and used for the selection of transfected ES cells in the tissue culture. The selection cassette was followed by a synthetic fragment introducing the modification. Precisely, the synthetic fragment comprises a 5’ part of Ex 14 (now Ex 14A), the artificial intron including the first LoxP site, mini Ex 14B (a short, 3’ part of Ex14 with size of 14 bp) and the second LoxP site 64 nucleotides (nt) downstream of Ex 14B. Accurate assembling of all foreigner fragments inserted into the standard cloning vectors was confirmed by complete sequencing.

The selected ES clones were PCR screened for presence of LoxP sites and artificial introns. In addition, for confirmation of correct homologous recombination with Dsg2 locus on the LA, a Southern blot analysis was performed in parallel).

A correctly targeted ES cells clones were injected into blastocysts to produce chimeric mice that transmitted the modified allele though the germ line. An animal, heterozygous for the targeted allele was bred with an ubiquitous Flippase (Flp) recombinase transgenic mice to ultimately produce animals that had deleted Neo cassette, preserving LoxP sites flanking the mini Ex 14B. Animals with correctly recombined alleles were intercrossed with Vil1-Cre transgenic mice constitutively expressing Cre recombinase in villus and crypt epithelial cells of the small and large intestines (B6.Cg-Tg(Vil1-cre)1000Gum/J; Jackson Laboratories). As a result, an Ex 14B was cut off, PSC was gathered and truncated form of Dsg2 protein, similar to the one detected in humans, was generated.

### Detection of homologous recombination in embryonic stem (ES) cells

To modify the Dsg2 locus in a conditional manner, C57BL/6N-derived embryonic stem (ES) cells were transfected with a K049.6 TV targeting vector. Thus, heterozygous cells containing both a targeted (TG) allele carrying the Dsg2 mutation and a WT allele were generated (supplemental figure 1). Prior to electroporation of ES cells, the targeting vector was linearized by digestion with NotI and SalI restriction enzymes. Since the K049.6 TV vector carries a Neo resistance gene, a selective antibiotic G418, at a final concentration of 0,2 mg/mL, was added to the maintaining cell culture medium resulting in accurate ES selection, where cells carrying Neo resistant construct were only growing. After 8 days of selection, 5 x 96 clones were picked and subjected to PCR analysis to verify a successful homologous recombination.

A screening PCR with a primer set that gives an amplicon only when a recombination event arises in the targeted locus, but not in any other random locus was performed. While the reverse primer (518LRPCR2) binds to the vector, within a region of the Neo cassette, the forward primer (K048.9) aligns specifically to the genome of the ES cells only, but not to the targeting vector, in a region just outside the homologous SA (supplemental table 3 and supplemental figure 3). Hence, if a signal is detected, a successful recombination between the targeting vector and the Dsg2 locus is validated.

To test the specificity of primer pair (518LRPCR2/K048.9), a control vector was assembled and consequently used as an internal positive control for screening PCR. The vector contains an elongated SA homology region, which holds 365 bp of extra genomic sequence that is not present in the targeting vector and is used as a template to which a K048.9 primer is selectively binding.

To confirm the presence of artificial intron and LoxP sites, a primer combination (K048.04/K048.05) (supplemental table 3) flanking this particular region was used. When the artificial intron and the LoxP site were present within the locus, the PCR product was longer (775 bp) compared to the much shorter WT fragment where the specific insertion was missing (446 bp).

Moreover, for confirmation of successful homologous recombination with the Dsg2 locus, a Southern blot was also performed.

### Southern blot analysis

DNA isolated from ES cell clones was digested with HindIII restriction enzyme (HindIII site locations are shown in supplemental figure 1 and consequently hybridized with a 3’ DNA external probe (LA probe) used for detection of the Dsg2 alleles (supplemental figure 1). The [*α-32P*] *dCTP DNA* external probe corresponds to a genomic sequence downstream of the incorporated mutation and it was generated using K048.16/ K048.17 primer pair (Supplemental table 3). Due to the size of the Neo cassette, the subdivided Ex 14 and the inserted in between an artificial intron, the TG allele is larger than the WT allele, hence, the size of the restriction fragment detected by the probes was 10.6 kb. In contrast, presence of a signal corresponding to 8.3 kb, confirmed the existence of a WT allele.

### Blastocyst injection and breeding

The positive ES cell clones were injected into blastocysts from grey C57BL/6N mice to obtain chimeras. The surviving blastocysts were then transferred into the uterus of pseudo-pregnant animals, i.e. CD-1 foster mice. To remove the FRT flanked Neo-resistance cassette from the TG locus, the newly born heterozygous transgenic mice were mated with Flp deleter stains, a homozygous mice expressing site-specific recombinase flippase (Flp). Upon this breeding program, Flp recognizes the FRT sequences surrounding the Neo cassette, leading to occurrence of a recombination event and complete excision of Neo cassette. Thus, a black offspring carrying the targeting construct with a deleted Neo cassette was established.

### Genotyping of offsprings from F1 generation

Genomic DNA extracted from tail or ear tissue was use as a template for a series of different PCRs verifying the genotype of first generation (F1) pups.

Flp-mediated deletion of the Neo cassette was verified by using K049.03/K049.04 primer combination (supplemental table 3 and supplemental figure 3), which align to the FRT sites remaining after successful deletion of the Neo cassette. For WT allele the anticipated amplicon is 383 bp in size, while for the targeted deleted Neo (TG del-neo) allele, a fragment with size of 502 bp is estimated.

To find out whether the offsprings still carry Neo cassette, a combination of primers (Neo.MP1/Neo.MP5/Neo.MP6) (supplemental table 3 and supplemental figure 3) was utilized. While Neo.MP1 binds simultaneously to a sequence from the Neo cassette and to a region within chromosome 3 of the mouse genome, Neo.MP5 and Neo.MP6 are unique and align to either of the above mentioned regions. Thus, PCR with Neo.MP1/Neo.MP6 primer pair results in an amplification of 512 bp-sized product, representing a region within the Neo cassette. Primers Neo.MP1/Neo.MP5 align to regions within the chromosome 3 and generate a product with size of 380 bp. The latter was used as an internal control. Moreover, screening for the presence of the artificial intron and the LoxP sites in the TG allele is achieved by K049.05/K049.2 primer pair (supplemental table 3). Primer K049.05 binds to the artificial intron and together with primer K049.02, which aligns to intron 14, result in a 471 bp PCR product for the TG allele. However, when WT allele is used as a template, no signal is detected. Additionally, a combination of primers SD24/SD25 (suppelmental table 3 and supplemental figure 3) was used to test for the existence of Flp recombinase. The latter was confirmed when a PCR fragment with size of 568 bp was observed.

### Transmission electron microscopy (TEM)

About 1 cm^2^ tissue was removed from colon of 5-7 days old mice and cut into 2 mm^2^ pieces for fixation in 2 % glutaraldehyde in PBS. The samples were washed in PBS and post fixed in 2 % osmium tetroxide. After successive hydration in 50 - 100 % ethanol series, the samples were cleared in propylene oxide and embedded in EPON 812 (SERVA Electrophoresis GmbH), and subsequently cured at 45 °C for 20 h and 60 °C for additional 24 h. The resulting blocks were trimmed and cut into 60 nm thin slices which were contrasted using methanolic uranyl acetate and aqueous lead nitrate. Images were captured using Libra transmission electron microscopy (Carl Zeiss NTS GmbH, Germany) equipped with a SSCCD camera system (TRS, Olympus, Tokyo, Japan).

### RNAseq

#### For mouse samples

Mouse intestine tissue samples were delivered to IMGM Laboratories GmbH in order to perform the analysis. Briefly, approximately 15-20 mg of mouse intestine tissue were used for RNA purification using the RNeasy Mini Kit (Qiagen, Hilden, Germany) according to the manufacturer’s instructions. The concentration and purity of the RNA was determined with the NanoDrop ND-1000 spectral photometer (Thermo Fisher Scientific, Waltham, MA, USA). The integrity of the RNA was subsequently evaluated using the 5300 Fragment Analyzer (Agilent Technologies, Santa Clara, CA, USA) using the high sensitivity total RNA 15nt Kit (Agilent Technologies, Santa Clara, CA, USA). Library preparation for gene expression analysis was performed with the NEBNext® Poly(A) mRNA magnetic isolation module and NEBNext® UltraTM II Directional mRNA kit (New England Biolabs, Ipswich, MA, USA) according to the manufacturer’s instructions with 1μg total RNA as input material. The quality of the RNA libraries was validated using the 2100 Bio Analyzer (Agilent Technologies, Santa Clara, CA, USA). Furthermore, all libraries were quantified using the highly sensitive fluorescent dye-based Qubit® ds DNA HS Assay Kit (Thermo Fisher Scientific, Waltham, MA, USA) and sequencing was performed using the illumina NovaSeq 6000 system (Illumina, San Diego, CA, USA).

#### For human samples

After ethical approval (142/16, 42/16, 113/13) RNA from patient samples was isolated using TRIZOL. RNA quality was checked using a 2100 Bioanalyzer with the RNA 6000 Nano kit (Agilent Technologies, Santa Clara, CA, USA). DNA libraries suitable for sequencing were prepared from 10 ng of DNase-treated total RNA, using SMARTer Stranded Total RNA-Seq Kit v2 - Pico Input Mammalian (Takara, Kusatsu, Japan). The fragmentation time was adjusted depending on the quality of the RNA input. After 12 PCR cycles, the size distribution of the barcoded DNA libraries was estimated ∼500 bp by electrophoresis on Agilent DNA HS Bioanalyzer microfluidic chips. Sequencing of pooled libraries, spiked with 5% PhiX control library, was performed at 25 million reads/sample in single-end mode with 75 nt read length on the NextSeq 500 platform (Illumina, San Diego, CA, USA) using High output sequencing kit. Demultiplexed FASTQ files were generated with bcl2fastq2 v2.20.0.422 (Illumina, San Diego, CA, USA).

To assure high sequence quality, Illumina reads were quality- and adapter-trimmed via Cutadapt^22^ version 1.16 using a cutoff Phred score of 20 in NextSeq mode, and reads without any remaining bases were discarded (command line parameters: --nextseq-trim=20 -m 1 -a AGATCGGAAGAGCACACGTCTGAACTCCAGTCAC). Processed reads were subsequently mapped to the human genome (GRCh38.p13 primary assembly and mitochondrion) using STAR^23^ v2.7.2b with default parameters based on RefSeq annotation version 109.20190905 for GRCh38.p13. Read counts on exon level summarized for each gene were generated using featureCounts v1.6.4 from the Subread package^24^. Multi-mapping and multi-overlapping reads were counted strand-specific and reversely stranded with a fractional count for each alignment and overlapping feature (command line parameters: -s 2 -t exon -M -O --fraction). The output count matrix was utilized to identify differentially expressed genes using DESeq2^25^ version 1.24.0. Read counts were normalized by DESeq2 and fold-change shrinkage was applied by setting the parameter “betaPrior=TRUE”. Differential expression of genes was assumed at an adjusted p-value (padj) after Benjamini-Hochberg correction < 0.05 and |log2FoldChange| ≥ 1.

### Quantitative reverse transcription PCR (qRT-PCR)

Total RNA was prepared from approximately 15-20 mg intestine tissue using RNeasy Plus Mini Kit (Qiagen, Hilden, Germany) according to the manufacturer’s protocol. cDNA was synthesized from 1000ng total RNA using SuperScript^TM^ II Reverse Transcriptase (Invitrogen, Waltham, MA, USA). Quantitative reverse transcription qRT-PCR was performed on a BioRad CFX96 Real-Time PCR detection system (Bio-Rad Laboratories Inc., Hercules, CA, U.S.) using SsoAdvanced^TM^ Universal SYBR Green Supermix (Bio-Rad Laboratories Inc., Hercules, CA, U.S.). Reactions were carried out in duplicate and nontemplate controls were included in every run. The following primer pairs were used: m*S100a8*: 5‘ - CCTTTGTCAGCTCCGTCTTCA - 3‘ (forward), 5‘ - TCCAGTTCAGACGGCATTGT - 3‘ (reverse); m*S100a9*: 5‘ - GCACAGTTGGCAACCTTTATG - 3‘ (forward), 5‘ - TGATTGTCCTGGTTTGTGTCC - 3‘ (reverse); m*Il17ra*: 5‘ – GCGCCGATCAAGAGAAACAT - 3‘ (forward), 5‘ – GAGTAGACGTCCAGACCTTCCTG - 3‘ (reverse); m*Gapdh*: 5‘ - AGGTCGGTGTGAACGGATTTG - 3‘ (forward), 5‘ – TGTAGACCATGTAGTTGAGGTCA - 3‘ (reverse). Primer sequences for *mS100a8* and *mS100a9* were published before^26^. Primer efficiencies were between 90% and 110%. Product specificity and amplicon size were controlled by gel analysis of the qPCR products. Relative expression of target gene levels were calculated according to the ΔΔCT method normalized to mGapdh levels and then wild-type mouse cohort using Bio-Rad CXF Maestro 2.3 software (Bio-Rad Laboratories Inc., Hercules, CA, U.S.).

## Results

### Identification of a CD patient with a rare mutation in the *DSG2* gene causing cytoplasmic truncation

In the summer of 1998, a then 19-year-old healthy female presented to the clinics and suffered from watery diarrhea, which later evolved into bloody diarrhea. Fever and pathogenic organisms in stool were absent. In September 1998, a colonoscopy was performed for the first time, which showed typical findings of CD including ileitis and colitis affecting the proximal colon (not shown). Few years later, in 2015, the patient presented with moderate colitis and a highly vascularized mucosa and in additional colonoscopies from 2017 and 2019 (figure 1A) the colitis was found to be progressive, developing into erythema and scarring changes of the mucosa. The histology was also typical for CD (figure 1B). The patient was first treated with mesalazine and glucocorticoids but in later years she was given immunosuppressants and biologic therapies because of the moderate to severe Crohn’s disease. At the age of 38 years she had prolonged pneumonia and enlarged lymph nodes in her inguinal areas. The T- and B-lymphocyte phenotyping showed slightly abnormal results. Thus, primary immunodeficiency (PID) was suspected and genetic testing was requested. Whole exome sequencing of the patient was performed but did not detect any clinically relevant variant to the patient’s phenotype. Incidental finding analysis according to the ACMG guidelines (https://www.ncbi.nlm.nih.gov/clinvar/docs/acgm/) revealed a heterozygous likely pathogenic variant in the *DSG2* gene: NM_001943.5 (DSG2):c.2315T>G (p.Leu772Ter) (figure 1C). The variant is absent in the Genome Aggregation Database (gnomAD) and has not yet been reported in the literature. The nonsense mutation is predicted to encode a truncated protein harbouring a termination codon in the cytoplasmic domain. Dsg2 immunostaining of a specimen from the patient’s colon obtained during resection revealed altered Dsg2 localization (figure 1B) in enterocytes of inflamed gut sections compared to other areas where Dsg2 expression from the remaining unaffected allele appeared normal (figure 1B). In follow-up, the patient has not been affected by prolonged or recurrent infections or adenopathy. Human DSG2 mutations are linked to heart-related diseases such as arrhythmogenic right ventricular cardiomyopathy (ARVC)^27^. However, the patient had no family history for IBD, ARVC or dilated cardiomyopathy. The family of the patient also underwent genetic screening. The mutation was detected in the healthy mother as well as in an aunt and two adult children of the patient diagnosed with celiac disease and psoriasis, respectively.

**Figure 1:**
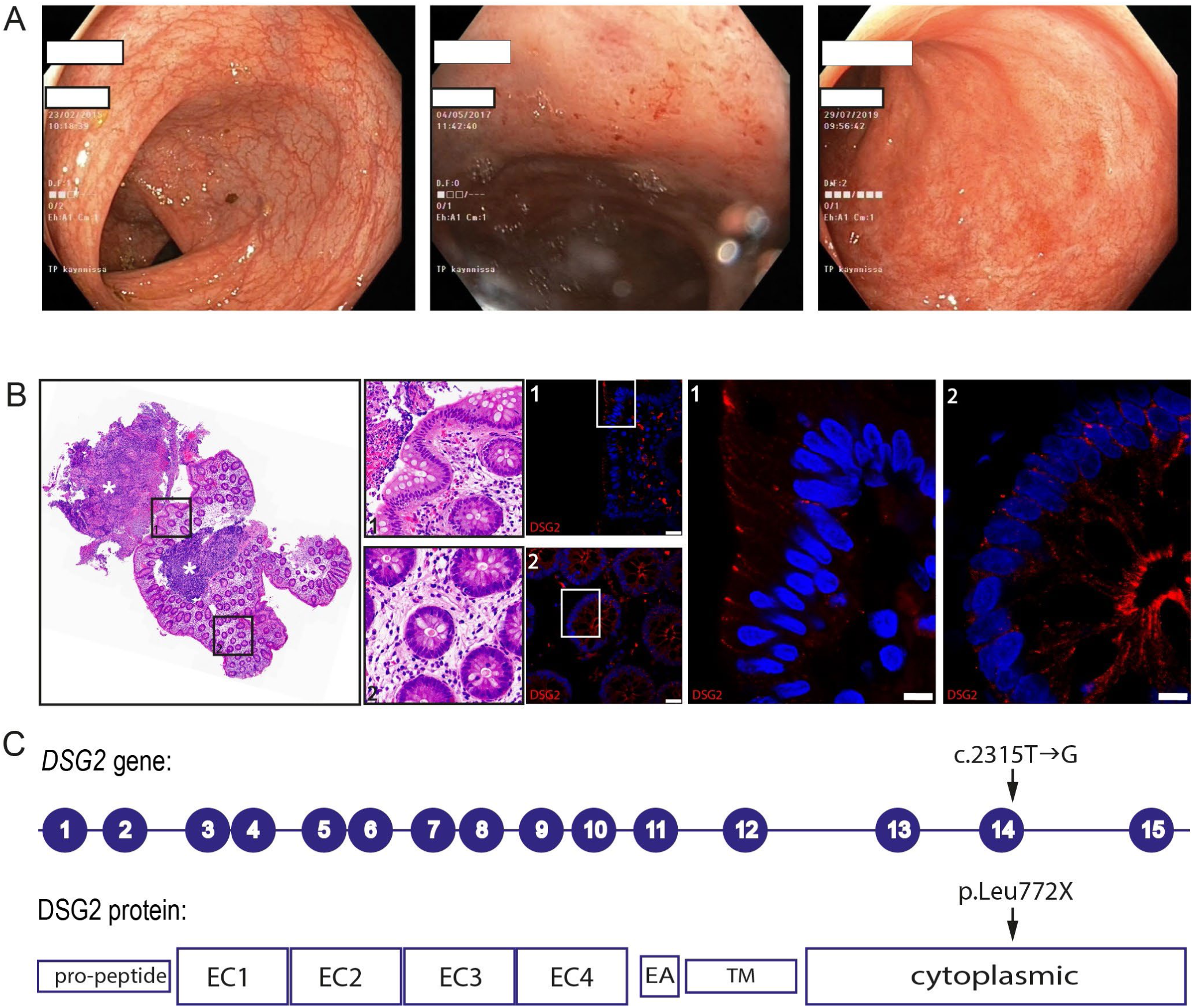
A patient harboring a pathogenic Dsg2 mutation presents with clinical and histological features suggestive of inflammatory bowel disease. **(A)** Images obtained during consecutive colonoscopies in 2015, 2017 and 2019, which reveal mucosal inflammation with ulcerations and progressive loss of vascularity. **(B)** Biopsy specimen taken from the ascending colon with hematoxylin and eosin staining (left panel, asterisks) shows inflammatory infiltrations, and immunostaining (right panel) for the extracellular domain of Dsg2 reveals fragmented Dsg2 localization. **(C)** Schematic illustrating the 15 coding exons of the human *DSG2* gene and the Dsg2 protein. The novel pathogenic mutation leads to a truncation of the terminal part of the cytoplasmic domain.

#### Dsg2 truncation in cultured enterocytes affects Dsg2 localization and mobility

Intrigued by the finding of a novel, likely pathogenic human mutation in the Dsg2 gene, we next investigated the molecular characteristics of the truncated Dsg2 gene *in vitro*. To assess how the cytoplasmic truncation of DSG2 affects it’ s expression on cellular level, a plasmid coding for the truncated DSG2, C-terminally tagged with an enhanced green fluorescent protein (eGFP), (hDsg2Leu772*-EGFP) was generated and transfected into DLD1-enterocytes lacking desmosomal proteins^15^. Western Blotting confirmed a lower band size for hDsg2Leu772* as compared to the wildtype protein (figure 2A). In cells reconstituted with wildytpe Dsg2, GFP immunofluorescence was very strong on cell-borders, while Dsg2Leu772* had a less prominent and rather dis-continuous localization at cellular membranes (figure 2B). The specificity of DSG2 expression was confirmed by co-immunostaining using an antibody against DSG2.

**Figure 2:**
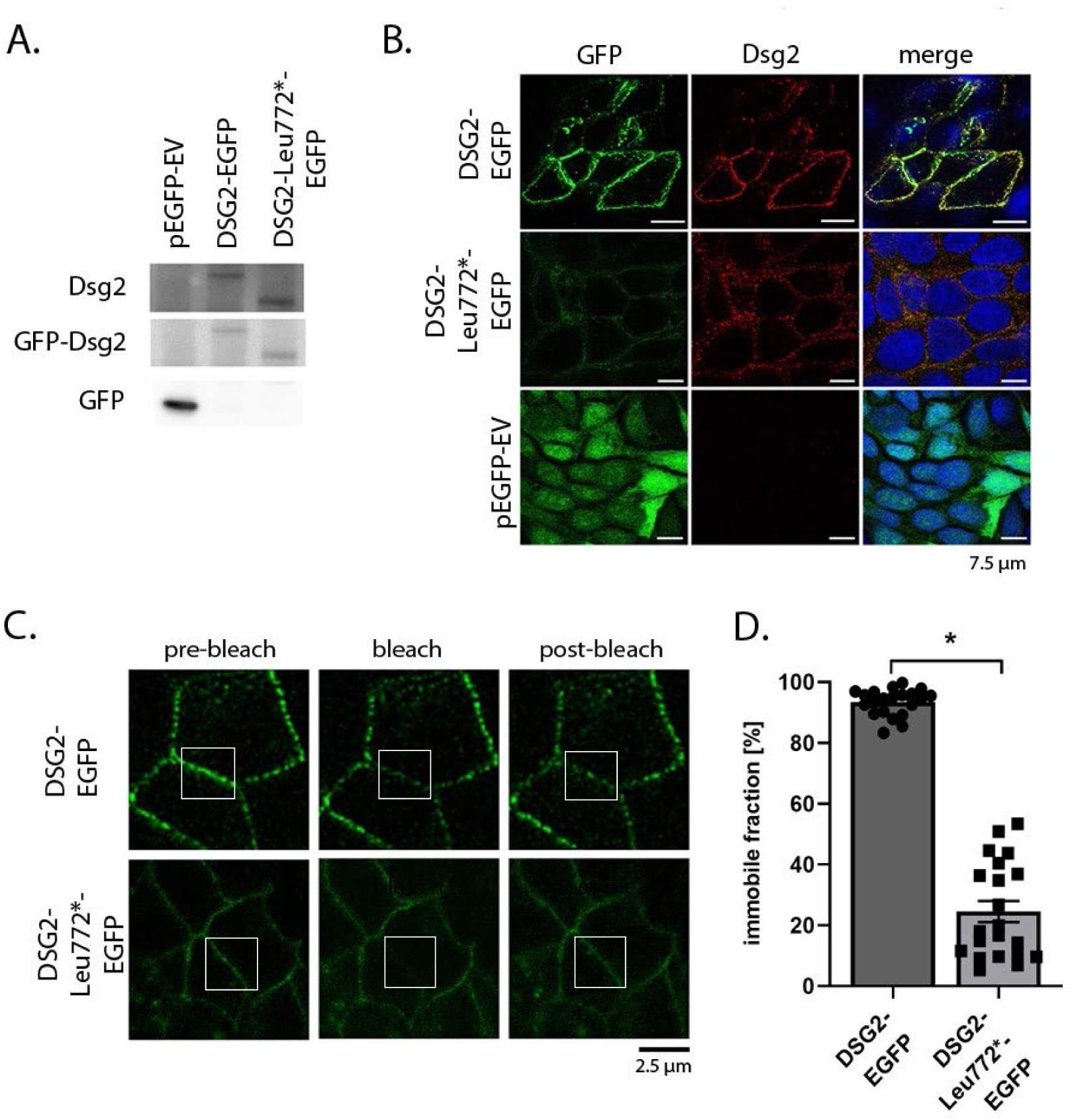
Truncation of Dsg2 affects Dsg2 localization and mobility at cell-cell borders. **(A)** Western blot analysis confirming the reconstitution of DLD-1 enterocytes defective for Dsg2 and Dsc2 (DLD-1 ΔDSC2 ΔDSG2) with the Dsg2 truncated form at Leucin 772 (DSG2Leu772*-EGFP) and Dsg2 (DSG2-EGFP) in contrast to the transfection with the control vector pEGFP-EV (n=4). **(B)** Immunofluorescence analysis after transfection of DLD-1 ΔDSC2 ΔDSG2 cells with the indicated plasmids. Co-staining with the Dsg2 autoantibody revealed successful and specific reconstitution of Dsg2. DSG2-EGFP and its truncated form is localized at cell borders. However, in cells that were reconstituted with DSG2Leu772*-EGFP a reduction of Dsg2 staining and localization on the cell borders were less continuous in comparison to the reconstitution with DSG2-EGFP. No specific staining was observed with the control vector pEGFP-EV. DAPI-staining proved the number and vitality of cells (n=3). **(C)** Representative images of FRAP of DLD-1 ΔDSC2 ΔDSG2 cells reconstituted with DSG2-EGFP and with DSG2Leu772*-EGFP revealing that the immobile fraction of truncated Dsg2 was significantly reduced in comparison to DSG2-EGFP. The white box highlights the bleached area. **(D)** Quantification of the average fraction of immobile DSG2Leu772*-EGFP and DSG2-EGFP is shown in Panel D (*p < 0.05; error bars represent SEM; N=4).

Next, the immobility of truncated Dsg2 was assessed by fluorescence recovery after photobleaching (FRAP) experiments, which revealed a less dynamic fraction of immobile proteins in Dsg2Leu772* transfected cells as compared to wildtype (figure 2C, D). These data demonstrate that Dsg2 cytoplasmic truncation severely altered Dsg2 localization and affects mobility.

#### The Dsg2 truncation mouse model is lethal and shows intestinal barrier defects

To clarify the pathogenicity of Dsg2 truncation for the IBD phenotype of the patient, we sought for a genetic mouse model. Complete genetic loss of Dsg2 in mice is known to be lethal early in development before formation of desmosomes and because Dsg2 is important for stem cell proliferation^28^. Therefore, we decided to establish the Dsg2 cytoplasmic truncation in an enterocyte-specific knock-in mouse model where Cre recombinase is driven by the villin promotor. The genetic mutation induces a premature stop codon and thus truncation of Dsg2 at the end of exon 14. The gene targeting strategy is summarized in figure 3A (figure 3A, detailed description is given in the Methods section). Mice expressing two copies of the floxed allele are further referred to as homozygous *Dsg2*_MUT_ or *Dsg2*_MUT_ mice. Western Blotting of colon tissue lysates revealed two weak Dsg2-immunoreactive bands in *Dsg2*_MUT_ mice (figure 3B), confirming the reduced expression of the truncated protein. Immunostaining against Dsg2, revealed fragmented Dsg2 expression in colon tissue from *Dsg2*_MUT_ mice (figure 3C), while wildtype Dsg2 shows a characteristic expression pattern in the epithelial layer. Homozygous *Dsg2*_MUT_ mice were indistinguishable from wild-type mice at birth and during the first days of life but did not thrive well in further course. A milk spot was present during the whole time, indicating regular suckling behaviour and nursing. However, homozygous *Dsg2*_MUT_ mice were significantly smaller than wild-type mice after five to ten days of life and had a reduced body weight (figure 3D-F). Kaplan-Meier curve showed that homozygous *Dsg2*_MUT_ mice but not wild-type mice died from seven to 14 days after age (figure 3G). Several gut segments in *Dsg2*_MUT_ mice displayed severe dilatation and gas bloat when compared to wild-type mice (figure 3H). Moreover, small intestine size, but not colon size, was significantly shorter in mutant mice which is indicative for intestinal barrier loss (figure 3I-J).

**Figure 3:**
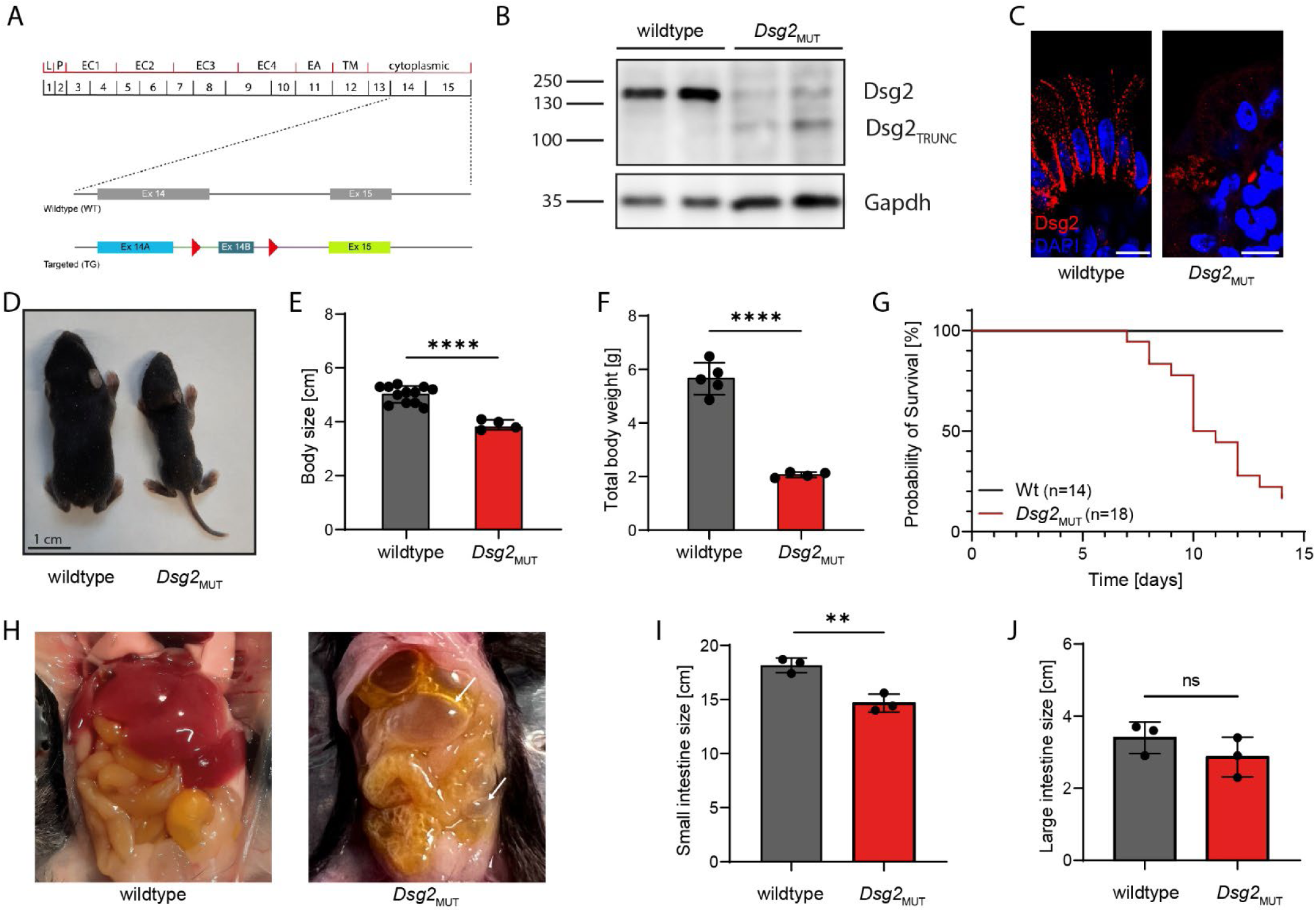
Characterization of *Dsg2* mutation in mice. **(A)** Schematic illustrating the strategy for generation of *Dsg2*_MUT_ mice. **(B)** Dsg2 protein level in murine colon samples was determined by western blotting using a Dsg2 antiserum. Dsg2 protein level in *Dsg2*_MUT_ mice is reduced and a truncated band is visible. **(C)** Immunofluorescence staining against Dsg2 in colon tissue reveals diminished Dsg2 immunostaining in enterocytes of *Dsg2*_MUT_ mice. Scale bar: 10 µm. **(D)** A photograph of 10 days old wild-type and *Dsg2*_MUT_ mice **(E, F)** Body size (E) and total body weight (F) of *Dsg2*_MUT_ mice are significantly reduced at 10 days of age compared to wild-type mice. **(G)** Kaplan-Meier Plot on the survival rate of *Dsg2*_MUT_ mice and wild-type litters. *Dsg2*_MUT_ mice start to die at postnatal day 7 and do not survive longer than day 14. **(H)** Representative photographs of 10 days old mice after sacrifice with opened peritoneum. Arrows indicate gas filled regions predominantly in distal small intestine and colon of *Dsg2*_MUT_ mice. **(I, J)** Length of small (I) but not large intestine (J) is smaller in *Dsg2*_MUT_ mice than in wild-type mice.

To further evaluate the properties of the intestinal barrier, we next performed a permeability assay on terminal ileum segments ex vivo. To this end, we measured the fluorescence intensity of fluorescein isothiocyanate (FITC)-conjugated dextran penetrating through the intestinal barrier over time (figure 4A, B). Intriguingly, dye transfer from the gut lumen was increased in *Dsg2*_MUT_ intestine segments, indicating a defective intestinal barrier. On morphological level, haematoxylin and eosin staining of colon segments demonstrated no gross mucosal damage but interstitial edema in *Dsg2*_MUT_ mice (figure 4C, arrows). On ultrastructure level, desmosomes in colon of *Dsg2*_MUT_ mice lack a proper keratin filament anchoring, appear asymmetric and display a larger intercellular space compared to wild-type littermates (figure 4D). Western Blot analysis and immunostaining in *Dsg2*_MUT_ mice demonstrated a significant up-regulation of pore-forming claudin 2 as well as of occludin whereas E-cadherin was not altered (figure 4E-G). Claudin 2 up-regulation was similar to that observed in patients with IBD^10^, reflecting a compromised barrier defect at the molecular level. Heterozygous mice were phenotypically inconspicuous during development. To test whether heterozygous mice show any barrier defect, we performed a FITC-dextran permeability assay on 14-days old mice (supplemental figure 4). Under basal conditions, heterozygous mice did not show a leaky intestinal barrier. However, when the mucosa was exposed to a cocktail of cytokines, *Dsg2*_MUT_^+/-^ mice showed a higher epithelial barrier permeability.

**Figure 4:**
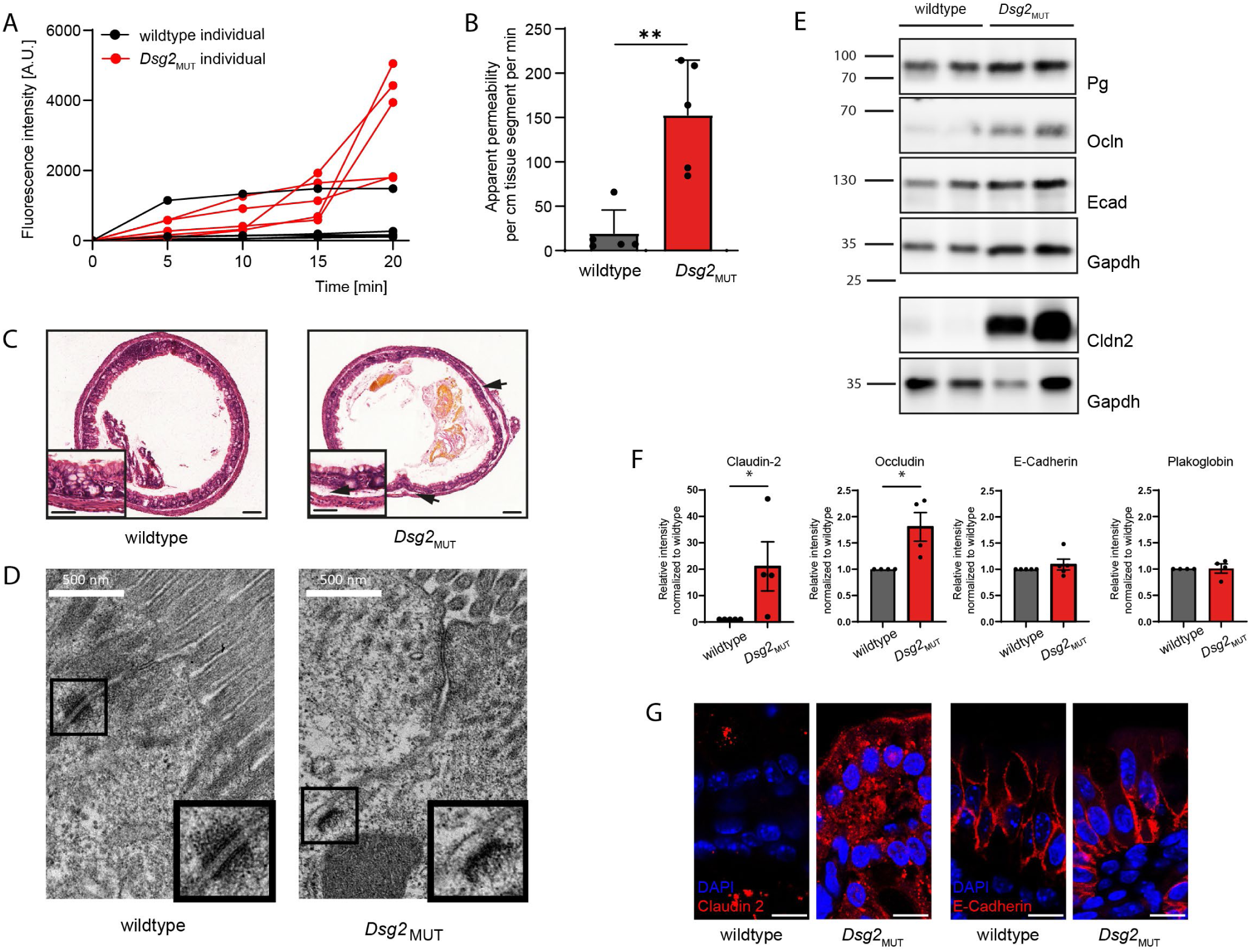
Cytoplasmic truncation of Dsg2 leads to intestinal barrier defect. **(A)** Levels of 4kDa FITC dextran fluorescent concentration in medium, in which a sealed, dye-filled intestinal segment was immersed, increased over time in *Dsg2*_MUT_ mice but not in wild-type mice, indicating a higher intestinal permeability *Dsg2*_MUT_ mice. **(B)** Slope of this increase in fluorescent intensity as shown in (A) corresponds to the apparent permeability (*p ˂ 0.05; n = 5). **(C)** Hematoxylin and eosin staining of colonic fragments. Notably, *Dsg2*_MUT_ mice exhibit no gross mucosal damage, but intestinal edema as indicated by arrows. Scale bar: 50 µm. **(D)** Electron microscopy revealed ultrastructural morphological alterations, including asymmetric appearance of the desmosomes and a widened intercellular space **(E)** Representative Western blotting analysis of protein expression of plakoglobin (Pg), occludin (Ocln), E-cadherin (Ecad) and claudin 2 (Cldn2) in colon tissue samples of *Dsg2*_MUT_ and wild-type mice. **(F)** Quantification of (E) The protein levels of claudin 2 and occludin are significantly increased in *Dsg2*_MUT_ mice (*p ˂ 0.05; n ≥ 4). **(G)** Accompanying immunostaining of colon tissue against claudin 2 and E-Cadherin in *Dsg2*_MUT_ mice and wild-type litters. Scale bar: 10 µm.

Taken together, these results demonstrate that homozygous cytoplasmic truncation of Dsg2 causes a severe lethal phenotype in mice with an intestinal epithelial barrier defect.

### Transcriptome analysis reveals an altered IL-17 signalling gene signature in both Dsg2_MUT_ mice and IBD patients

To assess the genetic profile of neonatal *Dsg2*_MUT_ mice and wildtype mice, we performed RNAseq analysis on colon tissue of 6 days old mice, which revealed a number of transcriptionally regulated genes (figure 5A). The RNAseq data set comprises 21 significantly upregulated and 28 significantly downregulated genes (adjusted p-value: padj < 0.05) in homozygous *Dsg2*_MUT_ mice, which are involved in inflammation and immune response (figure 5B). Intriguingly, S100a8 and S100a9 are among the most upregulated genes in *Dsg2*_MUT_ mice, which gene products are currently under investigation as candidate biomarker for IBD^29^. The upregulation of S100a8 and S100a9 was validated by qRT-PCR (figure 5C). Gene ontology analysis revealed a strong regulation of Il-17-mediated inflammatory response (figure 5D, Supplemental table 1), similar to what has been seen in the Dsg1^-/-^ mice epidermal transcriptome and psoriasis patients^26^. To assess the translational impact of the data, we compared the genetic profile of *Dsg2*_MUT_ mice with the transcriptome of a set of IBD patients, in whom a downregulation of Dsg2 was observed^11^. Mouse and human datasets have an overlap of 60 genes that were significantly upregulated and 7 significantly downregulated genes (figure 5E, supplemental table 2). Similar to *Dsg2*_MUT_ mice, IBD patients show a transcriptionally regulated Il-17/Il-23-mediated immune response (figure 5F), revealing a shared inflammatory signature between *Dsg2*_MUT_ mice and IBD patients (figure 5G).

**Figure 5:**
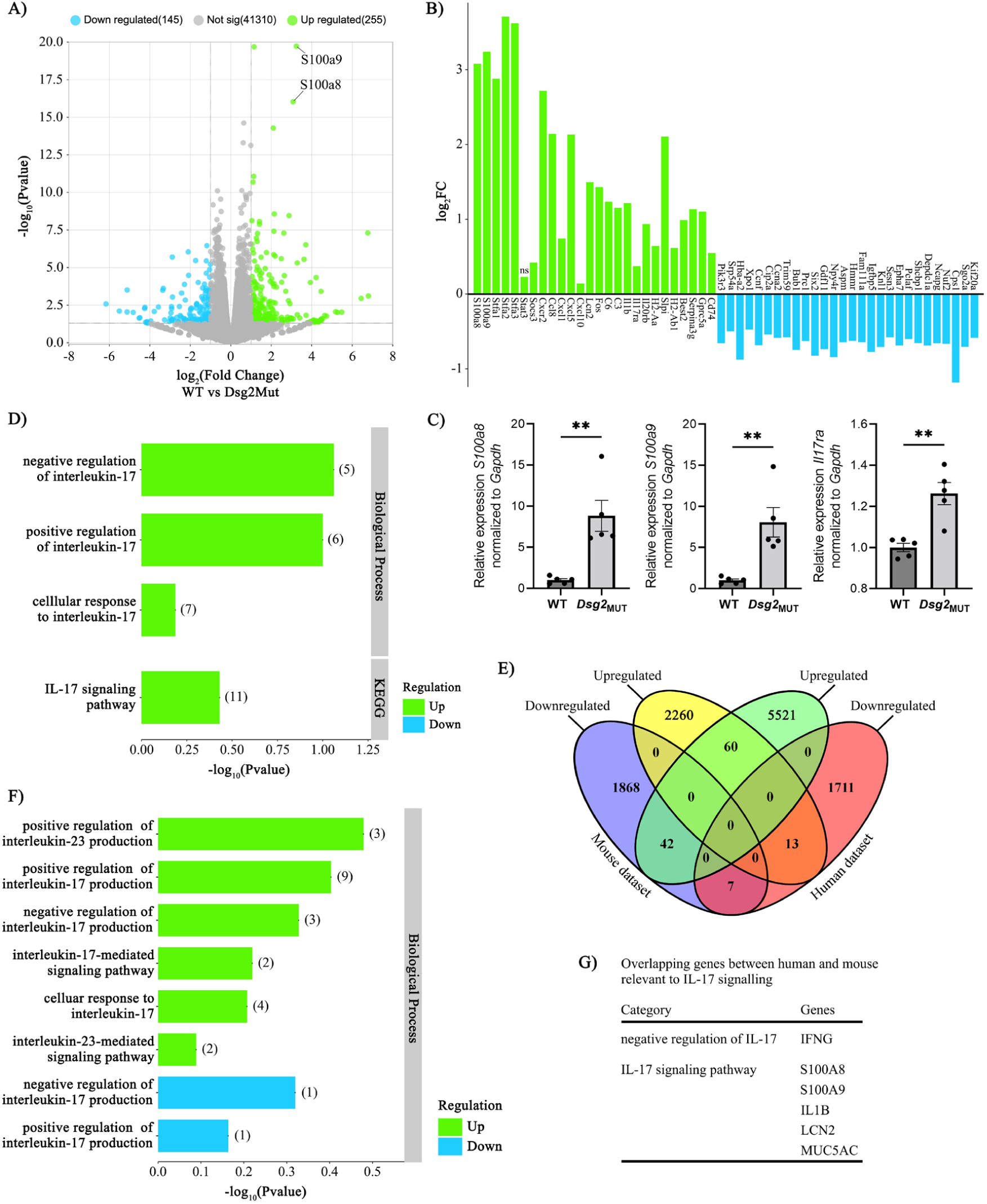
Differential gene expression analysis of RNAseq data from wild-type and *Dsg2*_MUT_ mice shows similarities with IBD patients harbouring a low Dsg2 expression. **(A)** Enhanced Volcano plot of upregulated and downregulated genes from neonatal mice colon. (n = 5/genotype). log_2_ FC ≥ 1 or ≤ −1 is considered significant. **(B)** mRNA expression levels for significantly upregulated or downregulated genes relevant for inflammation and immune response (FDR ≤ 0.05). **(C)** qRT-PCR data showing mRNA expression levels of S100A8, S100A9 and Il17ra in murine colon tissue. Gene expression of the cytokines was calculated according the ΔΔCT method and normalized to GAPDH (n = 5, **p < 0.005). **(D)** Go biological process gene ontology analysis depicting the number of upregulated genes involved in IL-17 regulation from *Dsg2*_MUT_ mice. Values in parentheses represent the number of genes contained within each category. **(E)** Venńs diagram representing the number of significant and non-significant genes with a log_2_ FC ≥ 1 or ≤ −1 from *Dsg2*_MUT_ mice and Dsg2-deficient human patients. Overlapping areas show the amount of common differentially regulated genes in both species. **(F)** Go biological process analysis depicting the number of upregulated genes involved in IL-17 and IL-23 regulation from Dsg2-deficient patients. **(G)** List of common genes differentially regulated in mouse and human relevant for IL-17 signalling.

## Discussion

The pathogenesis of IBD remains incompletely understood due to its complexity. It is well established that both intestinal epithelial barrier dysfunction and dysbalanced IL23/IL17 immune regulation play central roles for the pathogenesis of IBD^1–3 5^. Since only a fraction of patients has variants in the *IL23R* gene^30^, it is believed that immune dysfunction results from microbiota stimulating different sets of immune cells^7 31^. It can be envisaged that intestinal epithelial barrier defects occurring during the course of the disease aggravate this process. We demonstrate that genetic Dsg2 dysfunction found in a patient with CD caused skewing of the IL17 immune response in mice which is comparable to the transcriptome profile of IBD patients. This indicates that Dsg2 not only is crucial for intestinal barrier properties but in addition also serves as regulator of the immune system.

Here, we reported the first patient with IBD with a truncation of the cytoplasmic segment of Dsg2 which is required for cytoskeletal anchorage. Consistently, reconstitution of enterocytes with truncated Dsg2 affected junction localization and mobility of the protein. To prove pathogenic relevance, we established an enterocyte-specific knock-in model to induce cytoplasmic truncation of Dsg2 comparable to the IBD patient. Homozygous mice expressing truncated Dsg2 in colon did not thrive well and died within two weeks. Histology and immunostaining revealed mucosal defects and increased expression of claudin 2, and dye transfer showed impaired barrier properties in mice.

Noteworthy, these data do not demonstrate that the lethal outcome induced by homozygous Dsg2 truncation is caused by the barrier defect alone. Nevertheless, the findings are in line with a previous study reporting that ADAM17 deletion caused an inflammatory skin and bowel syndrome associated with alterations in Dsg2 turnover^32^. Importantly, ultrastructural alterations of desmosomes closely resembled the findings from CD patients^3 13^. Our data support the hypothesis that other factors were contributing to disease pathogenesis in the patient as well, especially since the patient’s mutation was heterozygous. ^33^. In general, it is believed that in most patients alterations in Dsg2 expression results from inflammation in response to inflammatory mediators such as TNF-α or reduced formation of growth factors including glial cell line-derived neurotrophic factor (GDNF)^10 11^. This study now reveals that impaired function of Dsg2 is sufficient to cause a severe intestinal barrier defect. Dsg2 truncation abrogated the cytoplasmic domain required for proper cytoskeletal anchorage resulting in enhanced mobility and reduced localization along cell junctions. The mechanisms by which Dsg2 contributes to intestinal epithelial integrity have been investigated during the last decade. Dsg2 is required for proper enterocyte adhesion and besides homophilic adhesion can also interact in heterophilic manner with Dsc2 and E-cadherin^34–36^. Moreover, Dsg2 can bind to the extracellular domain of EGFR, which is required for EGFR recruitment to cell junctions and determines EGFR function from cell proliferation towards cell adhesion, in which p38MAPK may also be involved^17 37^. More recently, Dsg2 was shown to sequester PI3-kinase and thereby preventing claudin 2 expression^38^. Dsg2 truncation mice also exhibited up-regulation of claudin 2 which most likely contributed to the intestinal barrier defect. In addition, intracellular Dsg2 cleavage has also been shown to sensitize cells to apoptosis^39 40^.

Completely new is the finding that Dsg2 regulates the IL17 immune response. This result is comparable to what has been shown for Dsg1 in the epidermis. A Dsg1-deficient mouse model, which was reported before to have a lethal epidermal barrier defect with histology resembling pemphigus foliaceus^41^, was shown to have a Th17-skewed inflammatory gene expression profile^26^. Interestingly, IL17-associated genes were dysregulated at E17.5 and thus before mice were born and exposed to microbiota. These data indicate that Dsg molecules can regulate expression of inflammatory molecules including S100A8 and S100A9, which were here found to be up-regulated in both Dsg2-truncation mice and CD patients and are known to be involved in IBD pathogenesis^29 42^. In this context it is important that both S100A8 and S100A9 were found very recently to interact with Dsg2 which provides a functional link between Dsg2 and regulation of inflammation^43^. Finally, it is interesting that the immune signature of Dsg1-deficient mice was to some extent comparable to psoriasis patients^26^. A relative of the patient from our study bearing the same Dsg2 mutation also suffered from psoriasis. Taken together, the fact that Dsg1 and Dsg2 have similar functions to regulate the IL23/IL17 immune response explain why novel therapy approaches targeting these molecules can be used to treat both IBD and psoriasis, although with different efficacy^5 44^. These data argue for a functional diversification of desmosomal cadherins during evolution. Desmosomes have started to diversify in parallel to transition of simple epithelial to complex stratified epithelia including development of new Dsg isoforms such as Dsg1 and Dsg3^45^. Thus, it appears that at least some functions of Dsg2, which is the only Dsg isoform expressed in simple epithelium, were transferred to Dsg1 expressed in superficial layers of stratified epithelia only.

The new mouse model will be useful to study IBD pathogenesis in context to immune system dysregulation in the future. Moreover, since Dsg2 dysfunction is required for intestinal mucosal integrity and immune regulation, future therapeutic strategies may also focus on stabilization of desmosome adhesion. In this context, a tandem peptide to crosslink Dsg2 in enterocytes was shown to protect against DSS-induced colitis in mice^38^ and also in a mouse model for the autoimmune disease pemphigus, which also is caused by desmoglein dysfunction^46^. Taken together, desmosomes appear to be involved in both regulation of cell adhesion and epithelial barrier properties and now also emerge as signalling hubs ^47^ controlling the immune response in different epithelia such as in the intestines and the epidermis.

## Founding and Acknowledgments

This work was supported by the DFG priority program SPP 1782 to JW and NS. We thank Sabine Mühlsimer, Frank Bodenstaff, Michelle Hermann and Cathleen Plietz for excellent technical assistance.

## Authors’ contributions and approval

DK and ESB together with DTE, JN, MH, NB performed the experiments. AKEH contributed to sample preparation. CSch provided reagents and instrumentation. DK, ESB, AGP and CS evaluated data. KSV and OK provided patient samples and data. JW and DK contributed to the conception and design of the study. DK, ESB, AGP, MYR, NS, JW interpreted data and wrote the manuscript. All authors contributed to the manuscript, read and approved the final version.

## Data Availability Statement

The raw data supporting the conclusions of this article will be made available by the authors, without undue reservation.

