## Supplemental Figures and Tables for "Dsg2 truncation causes a lethal barrier breakdown in mice"

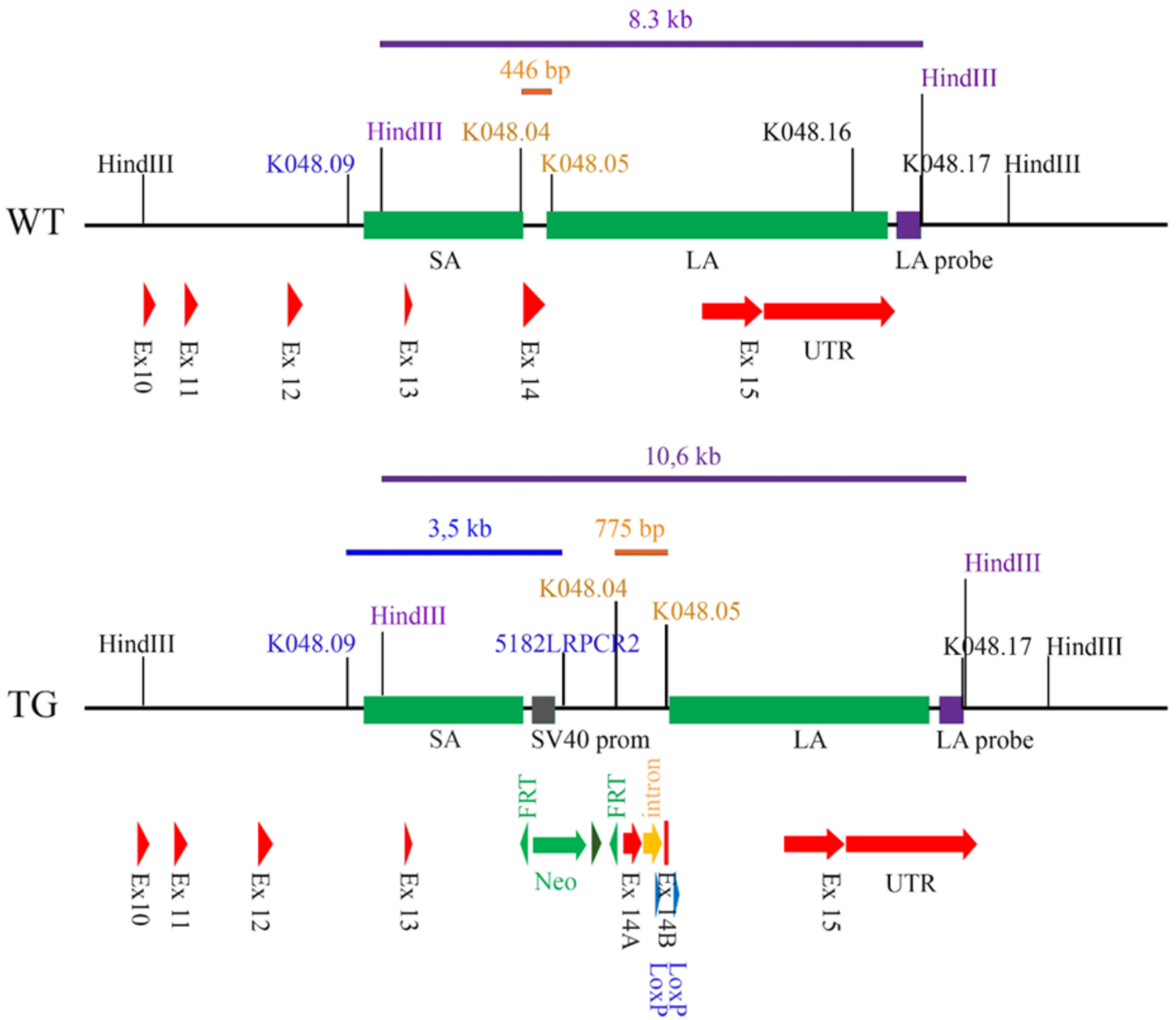


**Supplemental Figure 1. Schematic drawing of the different Dsg2 alleles carried by heterozygous ES cells**. The upper panel shows the WT allele, the lower panel the TG allele. Homologous regions (SA: short arm of homology; LA: long arm of homology) used for recombination is depicted as grey box. Dsg2 exon parts Ex 14A and Ex 14B are marked in red, separated by the artificial intron (orange), which contains the LoxP site (blue). For selection, an FRT-flanked neomycin resistance cassette (green) is inserted. The primers K048.9 and 5182LRPCR2 were used for PCR screening of recombined ES cells. The corresponding fragment size is indicated in blue. Primers used for confirmation of the presence of the intron including the LoxP sites are indicated in brown. The restriction enzyme HindIII, used for Southern blot analysis, and the corresponding external probe (LA-probe) and fragments are indicated in purple.

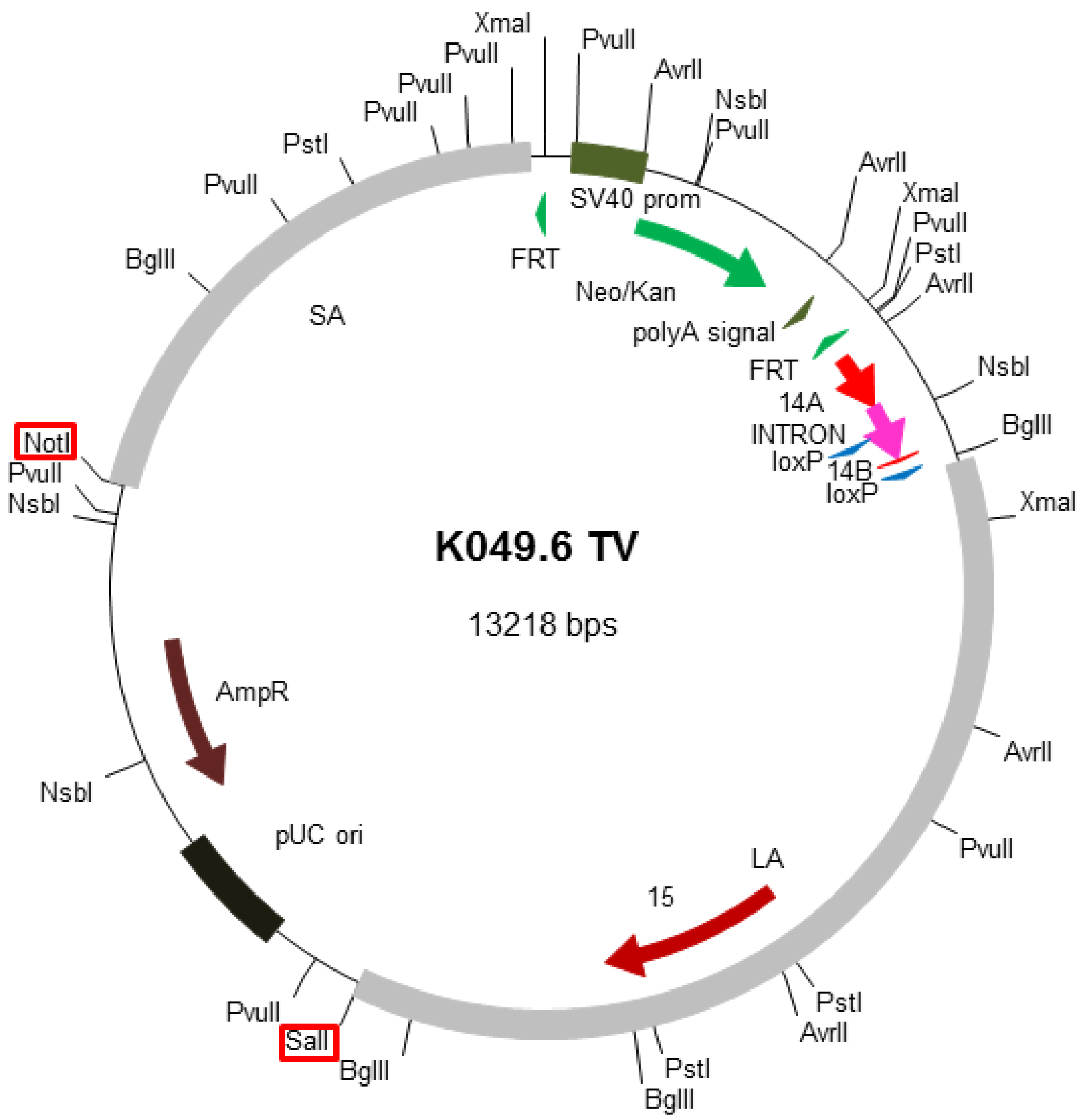


**Supplemental Figure 2: Targeting vector K046.6 TV**

Dsg2 exon parts 14A and 14B are marked in light red, separated by the artificial intron (pink), which contains the LoxP site (blue). For selection, an FRT-flanked Neo resistance cassette (green) is inserted. The LA has a length of 4.87 kb whereas the SA of homology extends for 2.72 kb (LA and SA; grey boxes). Restriction enzymes used for vector verification are indicated.

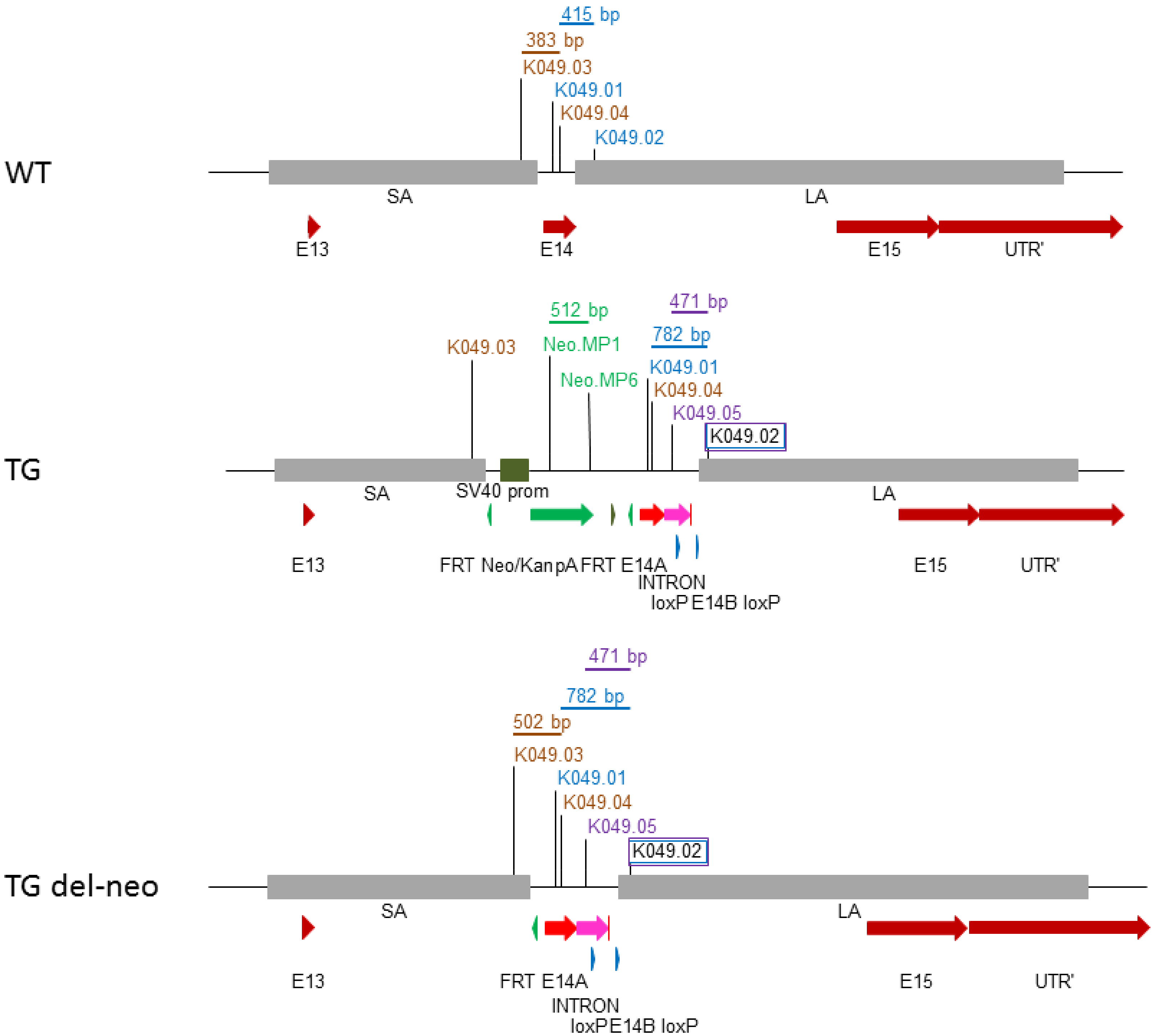


**Supplemental Figure 3**: **Schematic drawing of the WT locus and the targeted loci of Dsg2, with (TG) and without (TG del-neo) the presence of Neo resistance cassette**.
Dsg2 exon parts Ex 14A and Ex 14B are marked in light red, separated by the artificial intron (pink), which contains the LoxP site (blue). The primers used for genotyping and the corresponding amplicon sizes (brown: del-neo PCR, green: neomycin PCR, blue and purple: LoxP PCRs) are mapped on the different loci.

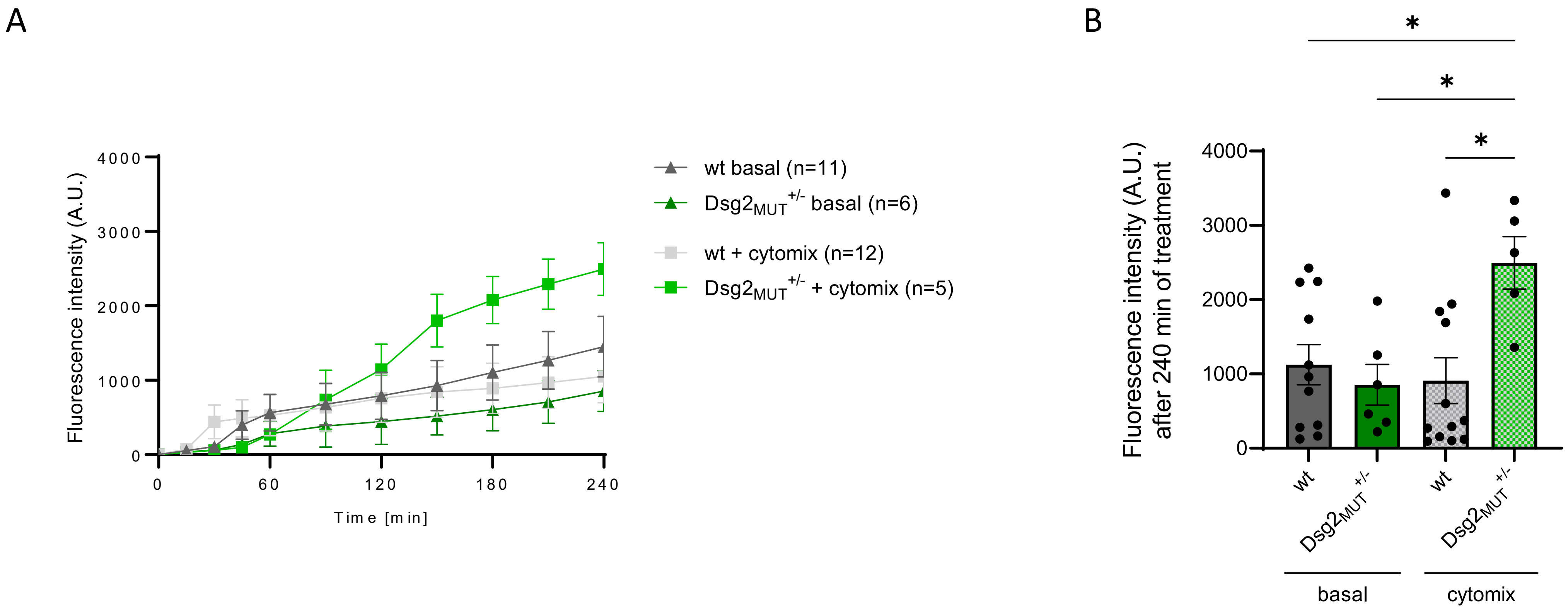


**Supplemental Figure 4: Intestinal permeability in terminal ileum segments of heterozygous *Dsg2*$_{MUT}$ mice (*Dsg2*$_{MUT}^{+/-}$) increases after mice were challenged with cytokine exposure.** Terminal ileum intestinal segments collected from 14 days old mice were filled with 2 µg/ml 4kDa FITC-dextran (basal) or 2 µg/ml 4kDa FITC-dextran substituted with TNFα, IFNγ and Il-1b (cytomix) and immersed into medium for a duration of 4 hours. **(A)** Medium was collected over the time-course at indicated time points and FITC-dextran intensity was photospectrometrically read out. **(B)** After 4 hours, 4kDa FITC-dextran fluorescence intensity in the medium collected from cytomix-treated heterozygous Dsg2$_{MUT}$ mice (Dsg2$_{MUT}$ $^{+/-}$) was significantly higher than in wildtype and under basal conditions. One-way ANOVA ($P<0.05$), followed by multiple comparisons test (Turkey-Test, alpha = 0.05). A Grubb's outlier test was performed for each group and one data point in the wt basal group (Fluorescence intensity = 5027) was identified and removed before statistical analysis.

**Supplementary Table 1**

| | *pValue* | *FDR* |
|---|---|---|
| ***Negative regulation interleukin-17*** | | |
| V-set immunoregulatory receptor (Vsir) | 0.030356939 | 0.236585343 |
| arginase type II (Arg2) | 0.415993 | 0.781048 |
| interferon gamma (Ifng) | 0.083232 | - |
| Interleukin-12b (Il12b) | 0.28306 | 0.677772 |
| tumor suppressor 2, mitochondrial calcium regulator (Tusc2) | 0.369351 | 0.749816 |
| ***Positive regulation interleukin-17*** | | |
| Interleukin-12b (Il12b) | 0.28306 | 0.677772 |
| Interleukin-18 (Il18) | 0.011149 | 0.133856 |
| myeloid differentiation primary response gene 88 (Myd88) | 0.664958 | 0.903328 |
| oncostatin M (Osm) | 0.233009 | 0.630194 |
| solute carrier family 7 (cationic amino acid transporter, y+ system), member 5 (Slc7a5) | 0.032111 | 0.244523 |
| tyrosine kinase 2 (Tyk2) | 0.456559 | 0.803112 |
| ***Cellular response to interleukin-17*** | | |
| chemokine (C-X-C motif) ligand 1 (Cxcl1) | 0.031767 | 0.242957 |
| chemokine (C-X-C motif) ligand 10 (Cxcl10) | 0.573762 | 0.863656 |
| decapping mRNA 1B (Dcp1b) | 0.07894 | 0.393192 |
| Interleukin-17 receptor A (Il17ra) | 0.001651 | 0.038491 |
| TRAF3 interacting protein 2 (Traf3ip2) | 0.213769 | 0.610486 |
| signal transducer and activator of transcription 3 (Stat3) | 0.020511 | 0.190591 |
| suppressor of cytokine signaling 3 (Socs3) | 0.001973 | 0.043087 |
| ***IL-17 Signaling pathway*** | | |
| CCAAT/enhancer binding protein (C/EBP), beta (Cebpb) | 0.006188 | 0.091293 |
| S100 calcium binding protein A8 (calgranulin A) (S100a8) | 9.26E-17 | 5.95E-13 |
| S100 calcium binding protein A9 (calgranulin B) (S100a9) | 1.87E-20 | 1.99E-16 |
| TNF receptor-associated factor 5 (Traf5) | 0.290761 | 0.683929 |
| caspase 8 (Casp8) | 0.634428 | 0.890484 |
| inhibitor of kappaB kinase epsilon (Ikbke) | 0.219683 | 0.617172 |
| Interleukin-1b (Il1b) | 0.001556 | 0.036893 |
| Interleukin-17F (Il17f) | 0.779071 | - |
| lipocalin 2 (Lcn2) | 2.04E-06 | 0.000334 |
| matrix metallopeptidase 9 (Mmp9) | 0.754325 | 0.933922 |
| mucin 5, subtypes A and C, tracheobronchial/gastric (Muc5ac) | 0.521459 | - |

**Supplementary Table 2**

| **UPREGULATED** | | | | **DOWNREGULATED** |
|---|---|---|---|---|
| ADM2 | GIP | MIRLET7D | SNORD17 | DSC3 |
| ALKAL1 | GPR141 | MUC5AC | SULT1E1 | FAM43B |
| ARRDC5 | GPRC5A | NT5C1A | SYT8 | HSD17B6 |
| ATP6V1B1 | HEMGN | PHLDA2 | TACSTD2 | LY6G5C |
| CCL8 | IFNG | PLA2G2A | TEX22 | MCOLN3 |
| CHAC1 | IL1B | PRSS2 | TEX43 | SLC23A1 |
| CHIT1 | ITGAD | PTPRQ | TIGIT | TAF7L |
| CIB3 | KLRC3 | RASAL1 | TMEM249 | |
| CNFN | KMO | REG3G | TNFRSF17 | |
| CPLX3 | L1TD1 | S100A3 | TNFRSF8 | |
| CTLA4 | LCN12 | S100A8 | TPSAB1 | |
| CXCL3 | LCN2 | S100A9 | TRIM29 | |
| CXCL5 | LRG1 | SIM2 | | |
| CYTL1 | LYPD5 | SLPI | | |
| DPEP3 | MIR200A | SNORA30 | | |
| FOS | MIR27A | SNORA64 | | |

**Supplemental Table 1:** Genes contained within each of the GO enrichment categories presented in Fig. 5D and their corresponding pValue and FDR.

**Supplemental Table 2:** List of “common” genes in Venn´s diagram differentially regulated in both human and mouse from Fig. 5E.

### Supplemental Table 3

| Purpose/Aim | Primer name | Sequence 5'-> 3' |
|---|---|---|
| Generation of external probe for Southern Blot | K048.16 | TTGGCTGGTCATGAGTTG |
| Generation of external probe for Southern Blot | K048.17 | TGACCTCTGCCCACATAG |
| Screening PCR for successful homologous recombination in ES cells | K048.9 | ACTCGTGCACTCAAGCACATAC |
| Screening PCR for successful homologous recombination in ES cells | 5182LRPCR2 | GTTGTGCCCAGTCATAGCCGAATAG |
| Screening for the presence of artificial intron and LoxP sites in ES cell clones | K048.04 | GATCCCTGCAGGCACATCTTCCTGCCAAGTACAC |
| Screening for the presence of artificial intron and LoxP sites in ES cell clones | K048.05 | TCCCTCATCACCATCCATAG |
| Screening for presence of the Neo resistance cassette in F1 generation | Neo.MP1 | GCTGTGCTCCACGTTGTCAC |
| Screening for presence of the Neo resistance cassette in F1 generation | Neo.MP5 | GGAAAGCTGGGCTTGCATCTC |
| Screening for presence of the Neo resistance cassette in F1 generation | Neo.MP6 | GGAGCGGCGATACCGTAAAG |
| Screening for deletion of Neo resistance cassette in F1 generation | K049.03 | GTGTCCTCCTCTGGTTGTAG |
| Screening for deletion of Neo resistance cassette in F1 generation | K049.04 | TCTTCCCATCTTCCGTCAAC |
| Screening for the presence of artificial intron and LoxP sites in F1 generation | K049.01 | CAGCTCAGCTTCCGTTACC |
| Screening for the presence of artificial intron and LoxP sites in F1 generation | K049.02 | ACTCAGCCACTGGCTATTTC |
| Screening for the presence of artificial intron and LoxP sites in F1 generation | K049.05 | CCTCATCCGTGGTTAAGG |
| Screening for Flp recombinase allele in F1 generation | SD24 | CTAATGTTGTGGGAAATTGGAGC |
| Screening for Flp recombinase allele in F1 generation | SD25 | CTCGAGGATAACTTGTTTATTGC |
| Screening for correct genomic DNA sequence | mDsg2-gDNA-752bp-FW | CATGAGCTGTCTGAGGTTGAC |
| Screening for correct genomic DNA sequence | mDsg2-gDNA-752bp-REV | CTCAGCCACTGGCTATTTCAGTG |
| Screening for correct splicing using cDNA as a template | mDsg2-cDNA-608bp-FW | CTGGTGGCCGATCATGCAG |
| Screening for correct splicing using cDNA as a template | mDsg2-cDNA-608bp-REV | GGTTACCGCTTGCTCATAGTGC |
